# *Mycobacterium marinum* infection drives foam cell differentiation in zebrafish infection

**DOI:** 10.1101/319202

**Authors:** Matt D. Johansen, Joshua A. Kasparian, Elinor Hortle, Warwick J. Britton, Auriol C. Purdie, Stefan H. Oehlers

## Abstract

Host lipid metabolism is an important target for subversion by pathogenic mycobacteria such as *Mycobacterium tuberculosis*. The appearance of foam cells within the granuloma are well-characterised effects of chronic tuberculosis. The zebrafish-*Mycobacterium marinum* infection model recapitulates many aspects of human-M. tuberculosis infection and is used as a model to investigate the structural components of the mycobacterial granuloma. Here, we demonstrate that the zebrafish-*M. marinum* granuloma contains foam cells and that the transdifferentiation of macrophages into foam cells is driven by the mycobacterial ESX1 pathogenicity locus. This report demonstrates conservation of an important aspect of mycobacterial infection across species.

## 1. Introduction

Studies focused on the important human pathogen *Mycobacterium tuberculosis* (*Mtb*) have identified direct and indirect mycobacterial metabolism of host lipids, and the transformation of macrophages into foam cells, as important pathways in mycobacterial pathogenesis (Lovewell et al., 2016). During early stages of granuloma formation in Mtb infection, macrophage transformation into foamy macrophages is driven by the internalisation of low-density lipoprotein particles and the retention of esterified cholesterol in the form of lipid droplets (Cardona et al., 2009). Foamy macrophages have been identified as an important nutrient source sustaining *Mtb* during infection. Additionally, foamy macrophages have been implicated in inhibiting lymphocyte access to infected macrophages and the build-up of caseum at the centre of granulomas, resulting in granuloma breakdown (Dong et al., 2017; Pandey and Sassetti, 2008; Russell et al., 2009).

The zebrafish (*Danio rerio*) model is a powerful platform for the investigation of host-pathogen interactions. Due to their optical transparency, the zebrafish larval infection model facilitates multiday observation of mycobacterial pathogenesis during early stages of infection in real-time within a live vertebrate (Oehlers et al., 2015). Moreover, the larval zebrafish immune system offers the ability to observe bacterial interaction with an innate immune system, highly conserved with mammals (Matty *et al*. 2016). Use of the zebrafish-*M. marinum* infection model has uncovered key insights into the granuloma as a bacterially-driven haven supporting tuberculosis pathogenesis (Cronan et al., 2016; Davis and Ramakrishnan, 2009). However, little is known about conservation of altered host lipid metabolism as a conserved motif across mycobacterial infections.

## 2. Materials and Methods

### 2.1. Zebrafish handling

Adult zebrafish were housed at the Garvan Institute of Medical Research Biological Testing Facility (St Vincent’s Hospital AEC Approval 1511) and housed for infection experiments at the Centenary Institute (Sydney Local Health District AEC Approval 2016-037). Zebrafish embryos were obtained by natural spawning and embryos were raised at 28°C in E3 media. All experiments and procedures were completed in accordance with Sydney Local Health District animal ethics guidelines for zebrafish embryo research.

### 2.2. Infection of adult zebrafish

Adult zebrafish, between the ages of 3 months and 12 months, were infected with approximately 200 CFU *M. marinum* by intraperitoneal injection as previously described (Oehlers et al., 2015). Infected animals were recovered into 1 g/L salt water, then fed and monitored daily.

### 2.3. Histological processing of adult zebrafish for Oil Red O staining

Adult zebrafish were euthanized by anesthetic overdose at 2 weeks post-infection and fixed in 10% neutral buffered formalin for 2 days at 4°C. Fixed specimens were washed in PBS, 30% (w/v) sucrose, 50:50 solution of 30% sucrose and OCT, and a final wash in OCT before freezing. Tissue sections were cut at 20 μm on a Leica cryostat. Slides were re-fixed in 10% neutral buffered formalin, rinsed in propylene glycol, stained in 0.5% (w/v) Oil Red O dissolved in propylene glycol and counterstained with a 1% (w/v) solution of methylene blue.

### 2.4. Infection of zebrafish embryos

Zebrafish embryos were raised to 30-48 hours post fertilization and anaesthetised with 160 μg/mL tricaine (Sigma-Aldrich) prior to infection with approximately 200 CFU M strain *M. marinum* via caudal vein injection. A dose of approximately 1000 CFU ∆*ESX1 M. marinum* was injected to match parental strain burden at 5 dpi. Embryos were recovered into E3 supplemented with phenylthiourea.

### 2.5. Oil Red O staining

Oil Red O lipid staining on whole mount embryos was completed as previously described (Passeri et al., 2009). Briefly, embryos were individually imaged for bacterial distribution by fluorescent microscopy, fixed, and stained in Oil Red O (0.5% w/v in propylene glycol). Oil Red O staining intensity at sites of infection were quantified in ImageJ and calculated as the pixel density difference from uninfected tissue.

### 2.6. Imaging

Live zebrafish embryos were anaesthetized in tricaine and mounted in 3% methylcellulose for imaging on a Leica M205FA fluorescence stereomicroscope. Histological sections were imaged on a Leica DM6000B. Further image manipulation and/or bacterial quantification was carried out with Image J Software Version 1.51j.

### 2.7. Quantification of *M. marinum* burden by fluorescent pixel count

Infection burden was measured as the number of pixels in each embryo above background fluorescence in ImageJ (National Institutes of Health) and pixels counted using the ‘Analyse particles’ function (Matty et al., 2016).

### 2.8. Statistical analysis

Graphpad Prism was used to perform statistical testing by unpaired Student’s t-tests.

## 3. Results

### 3.1. Lipid accumulation is observed in adult zebrafish-*M. marinum* granulomas

Adult zebrafish infected with *M. marinum* form highly organized and caseous necrotic granulomas that recapitulate many important aspects of the human-*M. tuberculosis* granuloma (Cronan et al., 2016; Oehlers et al., 2015). To determine if lipids accumulated in these complex zebrafish-*M. marinum* granulomas, adult zebrafish (>3 months old) were infected by intraperitoneal injection of *M. marinum* and maintained for 2 weeks post-infection, a time point where the animals contain a mixture of cellular and necrotic granulomas (Oehlers et al., 2017; Oehlers et al., 2015). Strong Oil Red O staining was observed in the cellular layer of necrotic granulomas and weaker Oil Red O staining was observed throughout cellular granulomas (Figure 1A and 1B).

The observation that lipid accumulation in granulomas correlated with granuloma maturation is consistent with the hypothesized role for foam cells in promoting granuloma immunopathology (Russell et al., 2009). We hypothesize cellular granulomas with weak lipid accumulation mature into necrotic granulomas with heavier lipid accumulation. To investigate this hypothesis, we switched model system to the more experimentally amenable zebrafish embryo to characterize the genesis of foam cells through temporal and functional analyses.

**Figure 1:**
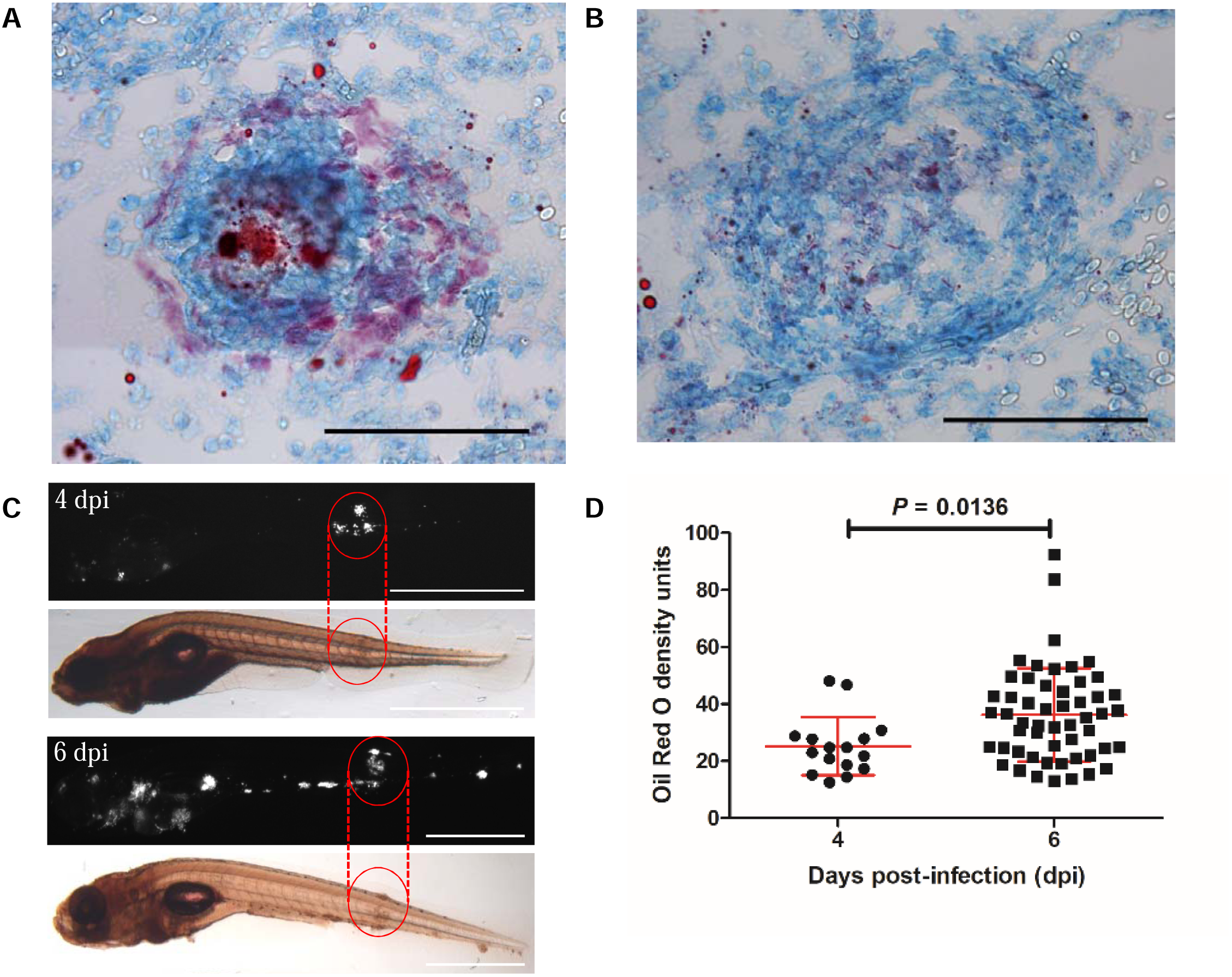
Lipid accumulation is present within the granulomas of zebrafish adults and embryos infected with *M. marinum*. *A,* Oil Red O staining of a necrotic granuloma from an adult zebrafish infected with *M. marinum* at 2 weeks post-infection. *B,* Oil Red O staining of a cellular granuloma from an adult zebrafish infected with *M. marinum* at 2 weeks post-infection. Note the heavier Oil Red O staining in the necrotic granuloma. Scale bars represent 100 μm. *C,* Representative images of Oil Red O staining density at 4 and 6 days post-infection. Red circles indicate areas of embryos with largest foci of infection. Scale bars represent 500 μm. *D,* Quantification of Oil Red O staining density of granulomas in zebrafish embryos at 4 and 6 days post-infection. Each data point represents the Oil Red O staining density of an individual granuloma. Error bars represent standard deviation, statistical tests were performed using an unpaired T-test.

### 3.2. Lipid accumulation is observed in zebrafish embryo-*M. marinum* granulomas

Zebrafish embryos lack an adaptive immune system and form simple *M. marinum*-containing granulomas primarily consisting of macrophages (Cronan et al., 2016). These simple granulomas have demonstrated remarkable immunopathology fidelity to more complicated adult zebrafish and mammalian granulomas (Cronan et al., 2016; Davis and Ramakrishnan, 2009; Oehlers et al., 2017; Oehlers et al., 2015). We investigated the temporal relationship of foam cell formation with known stages of granuloma maturation in the embryo model. Embryos were individually imaged for fluorescent mycobacterial distribution and stained with Oil Red O at 4 and 6 days post-infection (dpi) corresponding to pre-necrotic and necrotic granulomas, respectively. Granuloma lipid accumulation was first observed at 4 dpi and was increased in intensity at 6 dpi (Figures 1C and 1D).

### 3.3. Granuloma lipid accumulation is driven by mycobacterial pathogenesis

To establish if *M. marinum* drives foam cell transdifferentiation as a part of its granuloma-inducing pathogenesis program, we infected zebrafish embryos with ∆*ESX1 M. marinum*. The ∆ESX1 mutant strain is able to persist and multiply in zebrafish embryos, however it drives granuloma maturation to a much lesser degree than the parental strain (Davis and Ramakrishnan, 2009; Volkman et al., 2010). Compared to burden-matched embryos infected with WT *M. marinum*, ∆*ESX1*-infected larvae showed significantly less Oil Red O staining in mycobacterial granulomas (Figures 2A and 2B).

This experiment suggests foam cell genesis is driven by pathogenic mycobacteria as part of the ESX-1 virulence program. Taken with the circumstantial evidence that foam cells provide a niche for mycobacterial growth, and a haven from antibiotics and immune killing, this data suggests mycobacteria manipulate host lipid metabolism to drive transdifferentiation of permissive foam cells.

**Figure 2:**
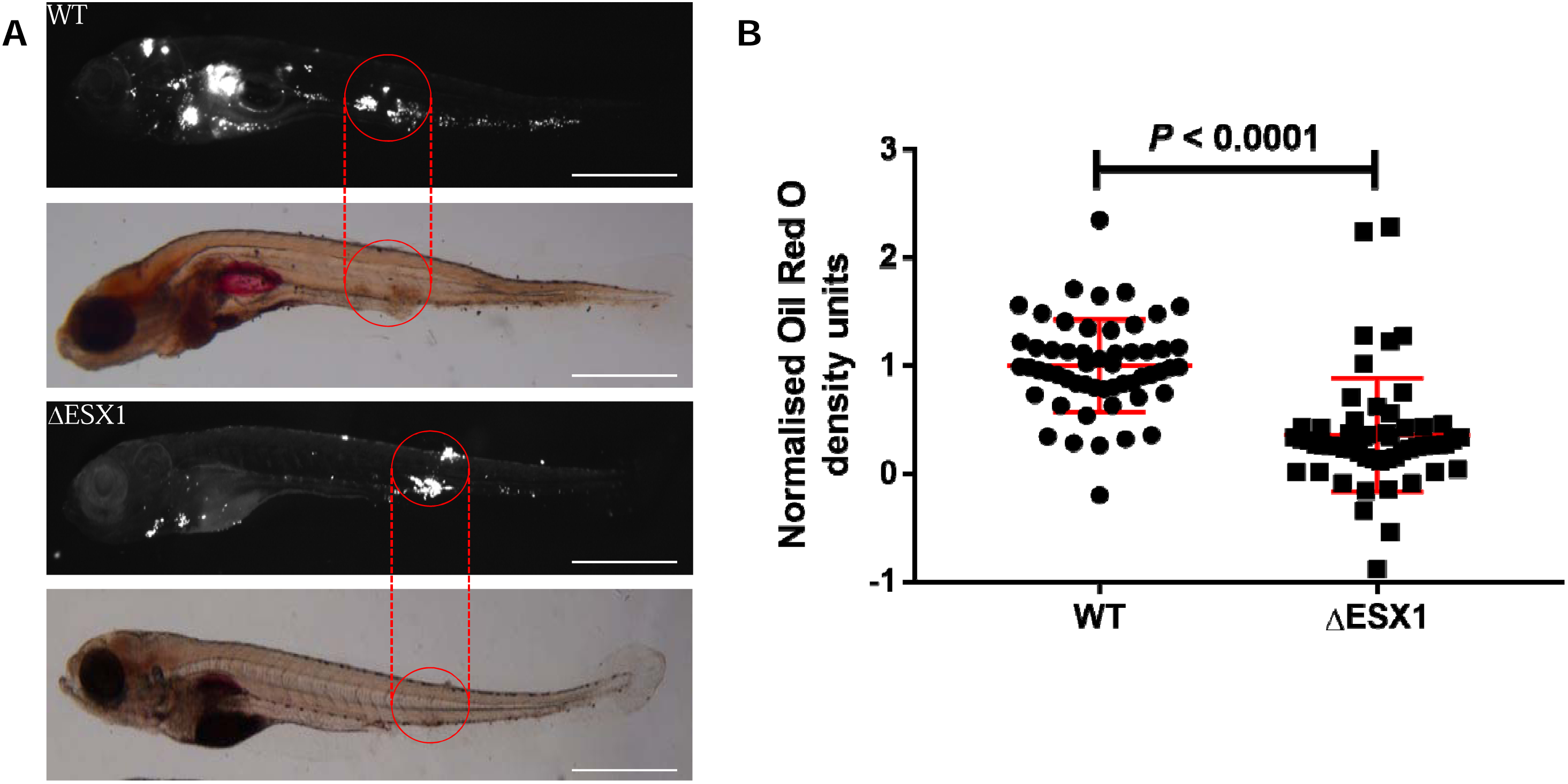
Granuloma lipid accumulation is driven by mycobacterial pathogenicity locus ESX1. *A,* Representative images of Oil Red O staining in zebrafish embryos infected with wild-type or ∆ESX1 *M. marinum* at 5 days post-infection. Red circles indicate areas of embryos with largest foci of infection. Scale bars represent 500 μm. *B,* Quantification of Oil Red O density units in embryos infected with wild-type or ∆ESX1 *M. marinum*. Each data point represents an individual granuloma. Error bars represent standard deviation. Analysis of Oil Red O density units was completed using an unpaired Student’s *t*-test.

## 4. Conclusion

This is the first report of mycobacterial infection-induced foam cells in the zebrafish model and recapitulates an important histological correlate of human TB disease in this important model organism. Our results show foam cell transdifferentiation is a correlate of granuloma maturation and that this accumulation of lipids in the granuloma is part of a highly conserved mycobacterial pathogenicity program. Our findings demonstrate the zebrafish-*M. marinum* platform will be applicable further mechanistic studies of infection-induced foam cell transdifferentation.

## Authorship contributions

Performed experiments: MDJ, JAK, EH, SHO. Supervised the study EH, WJB, ACP, SHO. Wrote the manuscript: JAK, SHO. All authors commented on the manuscript.

## Acknowledgements

Drs Anneliese Ashurst and Gayathri Nagalingam for training assistance with laboratory equipment; Dr Kristina Jahn and Sydney Cytometry for assistance with imaging equipment. Special thanks goes to the animal house staff at the Garvan Institute of Medical Research.

